# DataMed: Finding useful data across multiple biomedical data repositories

**DOI:** 10.1101/094888

**Authors:** L Ohno-Machado, SA Sansone, G Alter, I Fore, J Grethe, H Xu, Alejandra Gonzalez-Beltran, Philippe Rocca-Serra, Ergin Soysal, Nansu Zong, H Kim on behalf of the NIH BD2K bioCADDIE Consortium

## Abstract

The value of broadening searches for data across multiple repositories has been identified by the biomedical research community. As part of the NIH Big Data to Knowledge initiative, we work with an international community of researchers, service providers and knowledge experts to develop and test a data index and search engine, which are based on metadata extracted from various datasets in a range of repositories. DataMed is designed to be, for data, what PubMed has been for the scientific literature. DataMed supports Findability and Accessibility of datasets. These characteristics - along with Interoperability and Reusability - compose the four FAIR principles to facilitate knowledge discovery in today’s big data-intensive science landscape.

---

Biomedical research has always been a data intensive endeavor, but the amount of information was much more manageable just a decade ago. Today’s researchers not only have to stay abreast of the latest publications in their fields, but they also increasingly need to use existing data to help generate or test hypotheses *in-silico*, compare their data against reference or benchmark data, and contribute their own data to various “commons” to help health sciences move faster and be more easily reproducible^1,2^.

Data, software, and systems (e.g., analytical pipelines) are essential components of the ecosystem of contemporary biomedical and behavioral research. There are numerous databases focused, for example, on different communities, types of data, or types of research. While these may be indexed and searchable, these indexes usually do not interconnect, making it difficult to search for data across different research communities. Enabling broader focused searches for biomedical data was a key recommendation to the Director of the NIH by the Data and Informatics Working Group^3^. The work described here was funded to provide NIH with practical experience in fulfilling that recommendation.

## Starting the data discovery journey

The biomedical and healthCAre Data Discovery Index Ecosystem (bioCADDIE)^4^, is a Data Discovery Index Consortium funded by the NIH Big Data to Knowledge (BD2K) program^5^. The goal is to help users find data from sources which they would be unlikely to otherwise encounter, as PubMed does with the medical literature^6^. For example, biomedical researchers and clinicians don’t know the names of all journals that may have articles of interest. Even if they knew the journal names, it would be time consuming to search each resource separately. While searches within a certain journal web site could potentially be more detailed than those allowed in PubMed, they are not as useful if users do not get to those sites. PubMed, among other things, allows users to find articles in unfamiliar journals, and it makes sure that these journals meet certain quality criteria. Similarly, the bioCADDIE consortium is developed the search engine DataMed to help researchers find data of interest in a broad spectrum of high quality repositories.

A first prototype of DataMed is available at https://datamed.org. It performs searches on a shallow generic index that includes an initial set of 23 repositories and over 600k data sets, covering 10 data types. DataMed stores metadata generic enough to describe any dataset using a model we have called the DAta Tag Suite (DATS) (see working group 3(1) in Supplementary Box 1).

It is important to highlight some important differences between DataMed and broad harvesting of web-based data sets. DataMed is not the equivalent of Google Scholar^7^ or Microsoft Academic for data^8^.

a. bioCADDIE has developed criteria for data repository inclusion in DataMed (akin to journal inclusion in PubMed^9^) based on standards, interoperability, sustainability, overall quality, and user demand.
b. The DATS model, which is akin to the Journal Article Tag Suite (JATS) used in PubMed (NISO JATS Draft Version 1.1d3 April, 2015 http://jats.nlm.nih.gov/archiving/tag-library/1.1d3)^10^ enables submission of data for “ingestion” by DataMed (Figure 1). DATS has “core” metadata requirements that every data repository is expected to supply.
c. In addition to free text queries, DataMed provides a means to compose a structured query so that metadata specific to the life sciences can be better utilized.

**Figure 1.**
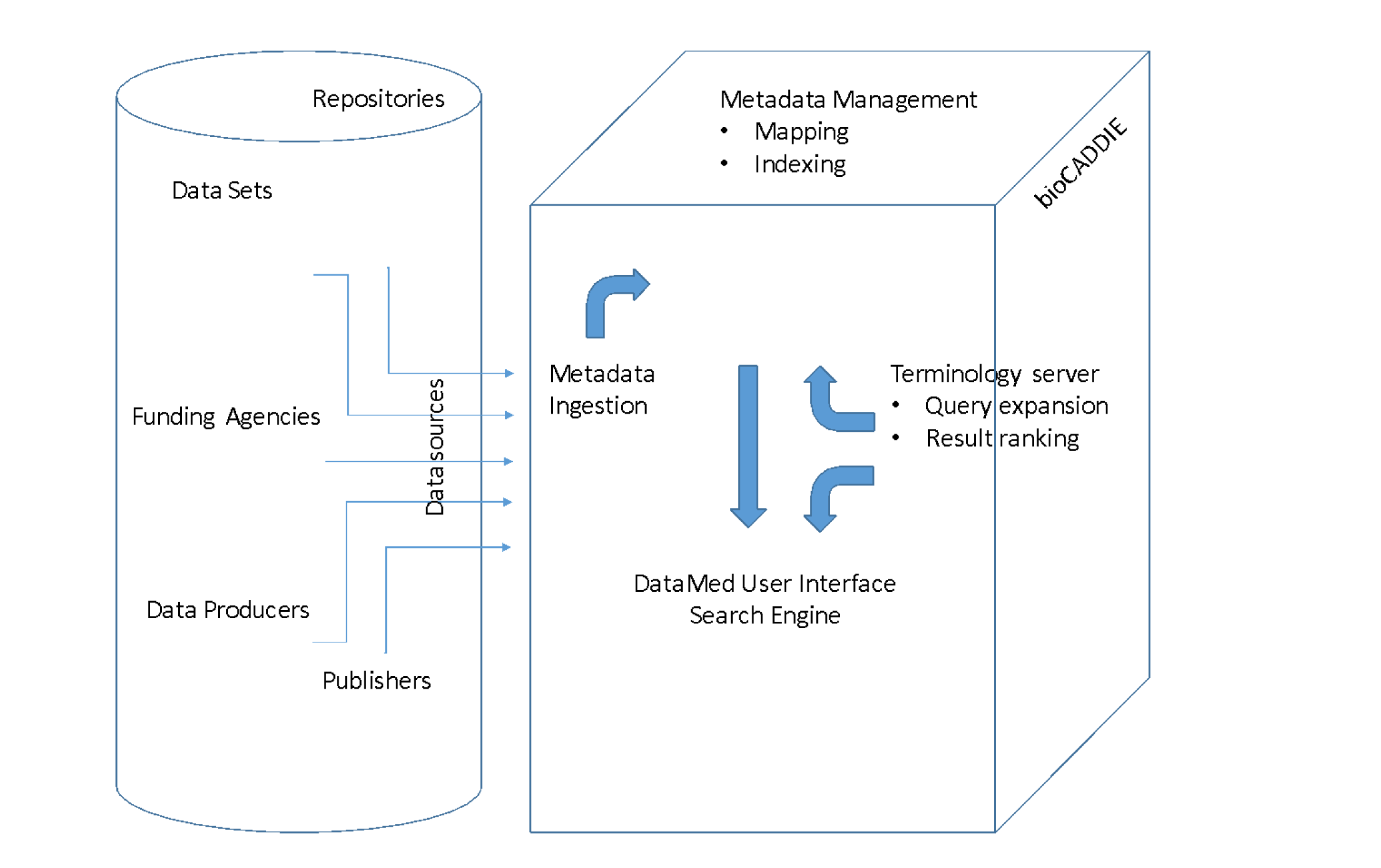
Data sources have various metadata specifications, which undergo “ingestion” into the common DATS model, whose metadata elements are used for indexing and DataMed searches. A terminology server is used to expand, transform, and standardize concepts used in metadata descriptions and in user queries.

DATS also facilitates the implementation of biomedical data discovery on other platforms in addition to DataMed. Building on prior attempts the use of structured data markup for web content is now reaching some maturity particularly in schema.org^11^. The bioCADDIE consortium worked with the scientific community on use cases for biomedical data search. DATS represents a platform independent model of the metadata necessary to support those use cases. The DATS model has been designed to be easily translated to and from *schema.org*^12^ (See working group 3(2) Supplementary Box 1). The mapping between DATS and schema.org and its biomedical extensions is expected to be seamless, with the potential benefit that data producers can expose their data assets to any web-based search tool. This opens the way for the emergence of new kinds of data-related web applications limited only by the ingenuity of their developers.

## Getting from point A to point Z: What is in between?

The specialized repositories that serve the needs of their specific communities, with their high level of specialization and more detailed metadata, are important components of the data discovery ecosystem. Although fine-grained metadata that are specific to certain areas are not yet indexed in DataMed, this search engine helps users find important data assets they are not aware of according to generic factors. We believe the amount of work to make the specialized data indexable by DataMed is very low - the better the quality of metadata in a repository, the easier it is to produce DATS metadata.

Looking more broadly at the ecosystem for data discoverability, a large community is participating in various aspects of bioCADDIE (Figure 2). Although the DataMed search engine interface is what users will interact with, we engaged community input on various components that operate behind the scenes to ensure that searches return the desired outputs. We formed Working Groups (WGs) and invited the research community to help us scope the project, select repositories, define indexing processes, develop a search engine and evaluate its results. The multiple WGs have included to date over 86 members from 56 institutions in the USA and the EU (see list of bioCADDIE collaborators and their affiliations). We expect to further broaden national and international participation as we get feedback from the community, develop new WGs, and continue the work on existing ones.

**Figure 2.**
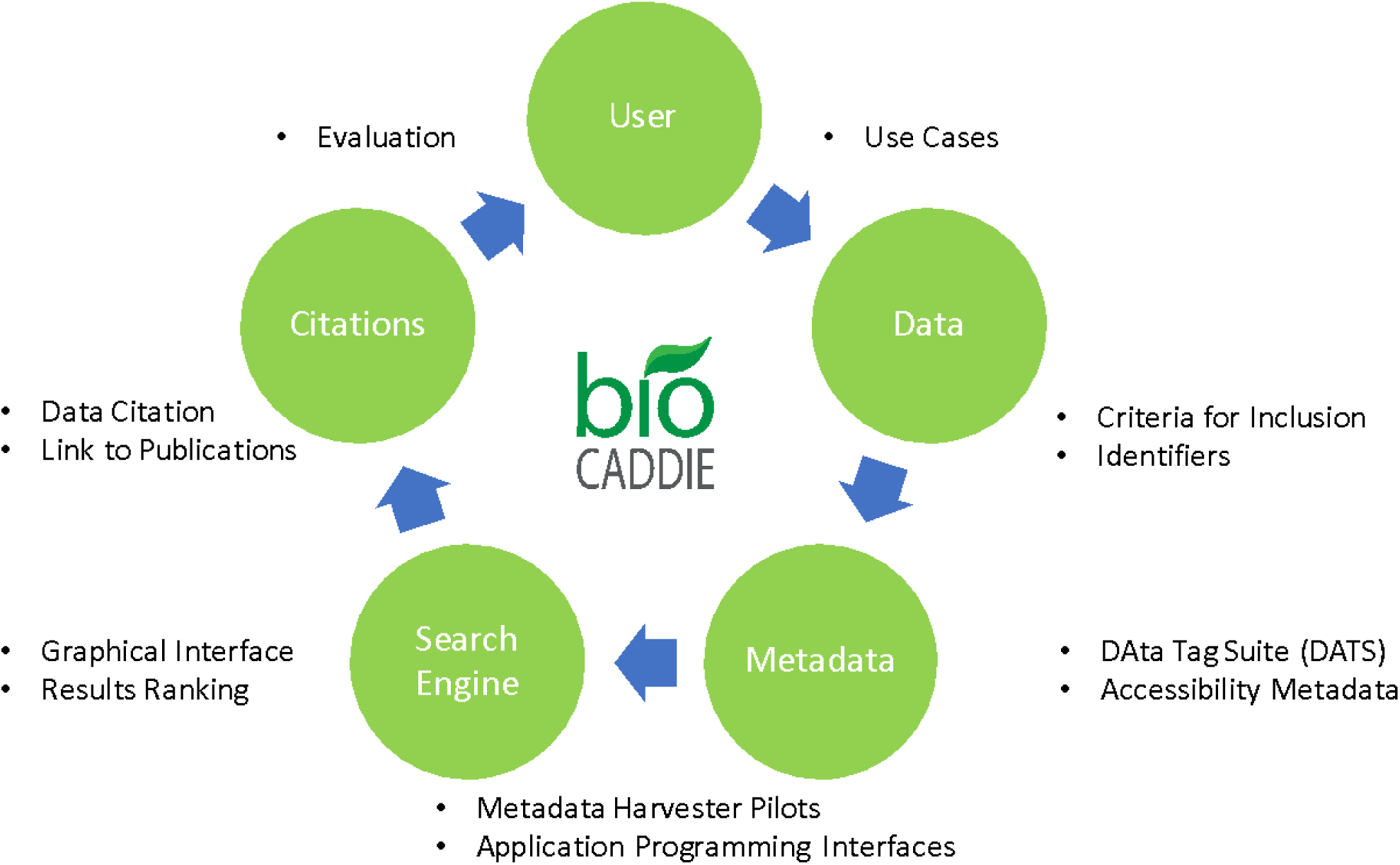
Community input to the Data Discovery Index Consortium. Working groups involved over 80 people from multiple institutions to scope the project via use cases, develop core metadata specifications, recommend identifier strategies, develop and test the search engine prototype, and discuss issues in data citation. Additionally, bioCADDIE funded external pilot projects for development of software that will be incorporated into the DataMed prototype.

## Organization

bioCADDIE’s activities leading to DataMed can be grouped into the following general areas:

### 1. Scope and Use Cases

Use cases helped define the appropriate boundaries and level of granularity for DataMed: which queries will be answered in full, which ones only partially, and which ones are out of scope. A workshop at the start of the project assembled researchers to discuss use cases. The current DataMed prototype is intended to point to data of interest by indicating the repository in which it is found, and providing some minimal information about the data so users can elect to follow the links to those repositories or move on to the next entry. Most use cases derived from community input (see working group 4 in Supplementary Box 1) are currently focused on finding data for a particular diagnosis and/or health condition (e.g., asthma) or data modality (e.g., fMRI).

### 2. Criteria for repository inclusion: standards, interoperability, sustainability

We established an initial set of criteria for a repository to be indexed by bioCADDIE (Supplementary Box 2). These criteria were inspired by those used by the National Library of Medicine in considering the indexing of a journal by PubMed^9^. An important consideration was to select data repositories that would help us evaluate search results (see working group 6 in Supplementary Box 1). We prioritized highly utilized repositories, because it would be hard to evaluate recall and precision if all repositories were unknown to users. Several new repositories are being added to the initial set currently available in DataMed.

### 3. The DATS model

The Metadata WG (see working group 3(1) in Supplementary Box 1) is a joint activity with the BD2K centers of excellence Metadata WG, and closely connected to other NIH initiatives (e.g., the BD2K Center for Data Annotation and Retrieval) and <u>[ELIXIR](https://www.elixir-europe.org/)</u> activities in Europe^13^. This group produced the DATS model, and its serialization describes the metadata needed for datasets to populate DataMed. DATS has a core and extended set of elements, to progressively accommodate more specialized data types. Like the JATS, the core DATS elements are generic and applicable to any type of datasets. The extended DATS includes an initial set of elements, some of which are specific for life, environmental and biomedical science domains and can be further extended as needed. The general concept fits with the idea of annotation profiles which allow a continuum from generic to specific annotation^14^. As with NISO-JATS the DATS model resides at the generic core of such a progression.

DATS has been designed to cover both (i) experimental datasets, which do not change after deposit in a repository, and (ii) datasets in reference knowledge bases describing dynamic concepts, such as “genes”, whose definition morphs over time. The core and the extended DATS entities are made available as machine readable JSON schemata, which will be annotated with schema.org elements.

The DATS model was developed given the following considerations:

- A variety of data discovery initiatives exists or are being developed; although they have different scope, use cases and approaches, the analysis of their metadata schemas is valuable. Several meta-models for representing metadata were reviewed (Supplementary Box 3) to determine essential items.
- Identification of the initial set of metadata elements was based on: (i) analyses of use cases (a *top-down* approach); and (ii) mapping of existing metadata schemas (a *bottom-up* approach). From the use cases, a set of ‘competency questions’ were derived; these defined the questions that we want DataMed to answer. The questions were abstracted, key concepts were highlighted, color-coded and binned in entities, attributes and values categories, to be easily matched with the results of the bottom-up approach.

The metadata schemas and models used in the mapping have been described in the <u>BioSharing Collection for bioCADDIE</u>^15^, which will be enriched progressively. Provided information includes: creators and maintainers; documentation, including URL where this is located; when the data set became available; version; source of metadata elements (e.g. XSD), including the URL where the model or schema was sourced.

#### Accessibility metadata

One of the strongest messages in the use cases was the need for information about data access^16^. Many types of biomedical data are restricted to protect confidential information about human subjects. Researchers want to know which data sets are readily available on the Internet and which ones require prior approval and other security clearances. We incorporated three dimensions in the core metadata (see working group 7 in Supplementary Box 1): authorization (Supplementary Note 1), authentication, and access type. Researchers also want to know which data can be accessed directly by machines through an application programming interface (API).

### 4. Identifiers and the data ingestion pipeline

#### Identifiers

We developed recommendations for the appropriate handling of identifiers for datasets and data repositories within DataMed (see working group 2 in Supplementary Box 1). At the most basic level, all data sets must be uniquely identified and web-resolvable. We will not mint identifiers for datasets, rather we will rely on the identifiers provided by the source. If the source does not have the capability to mint identifiers, we can suggest options for obtaining identifiers (e.g. issuing a DOI via Datacite^17^). In order to reuse identifiers provided by the data repository, a key set of features for a data repository supplied identifier are required. A identifier must be, (1) stable, (2) persistent for the life of the data repository, (3) unique within the data repository, and (4) resolvable (Supplementary Note 2) – i.e., a landing page must exist at the data repository that can accessed via embedding the identifier in a stable URL structure.

#### Indexing pipeline

The backend indexing pipeline for DataMed consists of a scalable architecture for ingesting, processing and indexing data. Metadata related to datasets are extracted and further enhanced by a document processing pipeline utilizing ApacheMQ^18^ and MongoDB^19^. Finalized metadata for a dataset is exported to an ElasticSearch endpoint that is then used by DataMed and can be used by external developers via its native RESTful API^20^ (Supplementary Note 3).

Operationally, to bring in a new data resource, a curator initially configures how the ingestion pipeline retrieves the metadata from a source (e.g. rsync^21^, OAI-PMH^22^, RESTful web service^23^) and provides the appropriate parameters. The ingestion pipeline then samples a number of records from the source for the curator so that an appropriate mapping can be made to the core metadata model described above. If the source has adopted a metadata standard that already exists within the pipeline (e.g. PDB XML^24^, GEO MINiML^25^, DATS (see working group 3(1) in Supplementary Box 1)) that mapping can be applied to the new source. However, if the source utilizes a different standard or a native API, a new mapping must be developed. After the initial mapping process a number of enhancement modules may be run to insert additional metadata into the dataset description document. Modules are being developed to enhance the pipeline, including an enhancer for semantic annotation (e.g. synonyms and super-classes)^26^ and one for citation altmetrics (i.e. the number of times a dataset has been cited)^27^. When a document has completed all processing steps it can be exported to a number of target endpoints via export modules. Currently, metadata records are exported to ElasticSearch whose native APIs are utilized by DataMed’s user interface.

### 5. Search engine prototype and usability testing

The Core technology Development Team (CDT)^28^ is developing the functioning prototype of DataMed with input from members of the community. The CDT has designed a modular architecture for the prototype backbone into which various projects can be integrated, creating a repository ingestion and indexing pipeline that maps to specifications provided by various working groups in the bioCADDIE community. The search engine interface provides all the functionality expected from a modern search application such as grouping the returning result via facets and saving/downloading search results. During search process, query input from user is processed to identify requested entities, and expanded with synonyms retrieved from a terminology server to enhance search results. This helps to prevent missing of datasets due to wording and terminological differences. Another important role for this it to host pilot projects, and integrate them into search interface seamlessly.

The CDT acquired user-specific input through the preliminary evaluation of DataMed by focus groups and by a limited number of end users (see working group 9 in Supplementary Box 1). Continuous evaluation will iteratively inform DataMed development. We designed and implemented the DataMed web application to provide a user-friendly interface that enables users to browse, search, obtain ranked results (see working group 8 in Supplementary Box 1) and in the near future get recommendations of related datasets tailored to their specific interests, preferences, and needs. A user-rating strategy is being discussed for future versions so that the application can learn from these ratings as well as from analyses of users behavior. The community may report issues on any aspect of DataMed through GitHub^29^ which provides a mechanism where they can be discussed and followed up in an open manner.

## Current status and next steps

The bioCADDIE consortium is continuing to engage stakeholders in the development and evaluation of metadata and tools. The quality of indexing is being optimized, as is the DataMed prototype search engine. Although highly utilized data repositories were the first to be included, there continues to be a need for indexing the “long tail” of science (i.e., smaller data sets that are being produced daily all over the world and may not be in repositories). Many new challenges will emerge, since it is unclear which data are going to be highly valued and how much assistance data producers will need in exposing their metadata for discovery. (Remember that PubMed relies on publishers, not individual authors.) Where to strike the right balance between quality and cost of maintaining the index from the viewpoints of data producers, data disseminators, and data consumers is still an open question. Nevertheless, we have to start somewhere, and exposing a resource early in its development (work on DataMed started in April 2015) helps us obtain valuable feedback from the user community to direct our next steps.

bioCADDIE and its products (core metadata specifications, criteria for repository inclusion, the DataMed search engine prototype) are intended to promote engagement and discussion around concepts that will last far beyond a particular grant or program. We invite everyone to join us in this journey to propel health sciences into a future where data are widely shared and easily discoverable and where discoveries relevant to human health are greatly accelerated.

## Acknowledgements

This project is funded by grant U24AI117966 from NIAID, NIH as part of the BD2K program.

The co-authors, which are the lead investigators and chairs/co-chairs of the core activities, thank all contributors to the bioCADDIE Consortium and list them here in alphabetical order within each activity group (each name appears only once even though many people participated in different activities)

Philip E. Bourne - NIH Associate Director for Data Science

## Steering Committee

<u>Alison Yao – NIAID, NIH</u>

<u>Dawei Lin – NIAID, NIH</u>

<u>Dianne Babski - National Library of Medicine, NIH</u>

George Komatsoulis - National Library of Medicine, <u>NIH</u>

<u>Heidi Sofia – NHGRI, NIH</u>

<u>Jennie Larkin – ADDS Office, NIH</u>

<u>Ron Mareolis – NIDDK, NIH</u>

## Metadata Working Group (WG)

<u>Allen Dearry – NIEHS, NIH</u>

<u>Carole Goble - The University of Manchester</u>

<u>Helen Berman - Rutgers</u>

<u>Jared Lyle - ICPSR, University of Michigan</u>

John Westbrook - Rutgers

<u>Kevin Read - NYU School of Medicine</u>

<u>Marc Twagirumukiza - W3C ScheMed WG</u>

<u>Marcelline Harris - University of Michigan</u>

<u>Mary Vardigan - ICPSR, University of Michigan</u>

<u>Matthew Brush - Oregon Health and Science University</u>

<u>Melissa Haendel - Monarch Initiative</u>

<u>Michael Braxenthaler - Roche</u>

<u>Michael Huerta - National Library of Medicine, NIH</u>

<u>Morris Swertz - CORBEL, BBMRI, ELXIR-NL and University Medical Center Groningen</u>

<u>Rai Winslow - John Hopkins University</u>

<u>Ram Gouripeddi - University of Utah</u>

<u>Shraddha Thakkar - NCTR FDA</u>

<u>Weida Tong - NCTR FDA</u>

## Accessibility Metadata WG

<u>Alex Kanous - University of Michigan</u>

<u>Anne-Marie Tassé - McGill University</u>

<u>Damon Davis - HealthData.gov</u>

<u>Frank Manion - University of Michigan</u>

<u>Jessica Scott - GlaxoSmithKline</u>

<u>Kendall Roark - Purdue University</u>

<u>Mark Phillips - McGill University</u>

<u>Reagan Moore - University of North Carolina</u>

## Identifiers WG

<u>Jo McEntyre - EMBL-EBI, ELIXIR EBI Node, Pub Med Central</u>

<u>Joan Starr - California Digital Library, DataCite</u>

<u>John Kunze - California Digital Library</u>

<u>Julie McMurry - Monarch Initiative</u>

<u>Merce Crosas - Data Science</u>

<u>Michel Dumontier - Center for Expanded Data Annotation and Retrieval, W3C HCLSIG</u>

<u>Stian Soiland-Reyes - University of Manchester</u>

<u>Tim Clark - Harvard Medical School, FORCE11 Data Citation Implementation Group</u>

## Use Cases and Testing Benchmarks WG

<u>Dave Kaufman - Arizona State University</u>

<u>Dina Demner-Fushman - National Library of Medicine, NIH</u>

<u>Ratna Rajesh Thangudu - BD2K Standards Coordinating Center</u>

<u>Steven Kleinstein - Yale School of Medicine</u>

<u>Thomas Radman - NIH</u>

<u>Todd Johnson - UTH</u>

<u>Trevor Cohen - UTH</u>

<u>Zhiyong Lu - National Library of Medicine, NIH</u>

March 2015 workshop use case contributors

## Ranking Search Results WG

<u>Chelsea Ju - University of California Los Angeles</u>

<u>Christina Kendziorski - University of Wisconsin Madison</u>

<u>Chun-Nan Hsu - UCSD</u>

Elmer Bernstam - UTH

<u>Gregory Gundersen - Icahn School of Medicine at Mount Sinai</u>

<u>Griffin Weber - Harvard Medical School</u>

<u>Hongfang Liu - Mayo Clinic</u>

Jim Zheng - UTH

<u>Sanda Harabagiu - University of Texas at Dallas</u>

<u>Vincent Kyi - University of California Los Angeles</u>

## Criteria for Repository Inclusion WG

<u>Amy Pienta - University of Michigan, ICPSR</u>

Elizabeth Bell – UCSD

<u>John Marcotte - ICPSR / University of Michigan</u>

<u>John Yates - The Scripps Research Institute</u>

<u>Kei Cheung - Yale University</u>

<u>Larry Clarke - National Cancer Institute (NCI)</u>

<u>Marek Grabowski - University of Virginia</u>

<u>Matthew McAuliffe - Center for Information Technology NIH</u>

<u>Neil McKenna - Baylor College of Medicine</u>

<u>Tanya Barrett - NCBI (GEO, BioSample, BioProject), GA4GH</u>

<u>Tim Clark - Harvard Medical School, FORCE11 Data Citation Implementation Group</u>

## Dataset Citation Metrics WG

<u>Daniel Mietchen - National Library of Medicine, NIH</u>

<u>Jennifer Lin - PLOS</u>

<u>Kristi Holmes - Northwestern University</u>

<u>Martin Fenner - DataCite</u>

<u>Michael Taylor - Elsevier</u>

<u>Rebecca Lawrence - F1000</u>

<u>Sunje Dallmeier-Tiessen - CERN</u>

<u>Trisha Cruse - DataONE, CDL</u>

## Core technology Development Team (CDT)

<u>Anupama Gururaj - UTH</u>

<u>Burak Ozyurt - UCSD</u>

<u>Claudiu Farcas - UCSD</u>

<u>Cui Tao - UTH</u>

<u>Deevakar Rogith - UTH</u>

<u>Ergin Soysal - UTH</u>

<u>Larry Lui - UCSD</u>

Mandana “Nina” Salimi - UTH

<u>Min Jiang - UTH</u>

<u>Muhammad Amith - UTH</u>

<u>Nansu Zong - UCSD</u>

Ngan T Nguyen-Le - UTH

Pratik Kumar Chaudhary - UTH

<u>Ruiling Liu - UTH</u>

<u>Saeid Pourneiati - UTH</u>

<u>Vidya Narayana - UTH</u>

<u>Xiao Dong - UTH</u>

<u>Xiaoling Chen - UTH</u>

<u>Yaoyun Zhang - UTH</u>

<u>Yueling Li - UCSD</u>

## Pilot project PIs and collaborators

Aditya Menon – National ICT Australia

Cathy Wu – University of Delaware

Cecilia Arighi – University of Delaware

Chris Mungall – Lawrence Berkeley National Laboratory

Dmitriy Dligach – Boston Children’s Hospital

Guergana Savova – Harvard University

Guoqian Jiang – Mayo Clinic

Harold Solbrig – Mayo Clinic

Hwanjo Yu – POSTECH, South Korea

Jaideep Vaidya – Rutgers

Jeeyae Choi – University of Wisconsin Milwaukee

Jina Huh – UCSD

Julio Facelli – University of Utah

Peter Rose – UCSD

Ricky Taira – University of California Los Angeles

Xiaoqian Jiang – UCSD

Zhaohui Qin – Emory University

## Supplements to bioCADDIE

OmicsDI

Eric Deutsch, ISB
Henning Hermjakob EMBL-EBI
Peipei Ping, UCLA

CountEverything

David Haussler, UC Santa Cruz
Ida Sim, UC San Francisco
Isaac Kohane, Harvard Medical School

## FORCE11

Tim Clark, Harvard Medical School

Maryann Martone, UC San Diego

## Administrative Supplements

Bert O’Malley, Baylor College of Medicine

Carolyn Mattingly, North Carolina State University Raleigh

George Hripcsak, Columbia University

Hongfang Lu, Mayo Clinic

Raymond Winslow, Johns Hopkins

Steven Kleinstein, Yale

Tor Wagner, University of Colorado

Trevor Cohen, UTH

